# Integration of ethologically defined anxiety-related behaviors in the ventral hippocampus

**DOI:** 10.64898/2026.01.24.701488

**Authors:** Kaizhen Li, Marios Kyprou, Stéphane Ciocchi

## Abstract

Anxiety is a complex emotional state that unfolds as structured sequences of risk assessment and exploratory behaviors enabling animals to evaluate potential threats in the environment. The ventral hippocampus (vH) regulates anxiety responses, yet the neuronal substrates orchestrating specific ethologically defined anxiety-related behaviors remain unclear. Here, we combined high-resolution 3D behavioral tracking with cell-type specific *in vivo* calcium imaging and optogenetic manipulations to dissect vH microcircuit dynamics during anxiety. We identified protected and unprotected stretch-attend postures (pSAP and uSAP), along with head dipping, as core anxiety-related risk-assessment behaviors organized along a spatial gradient in the elevated plus maze (EPM). Ventral hippocampal (vH) pyramidal neurons were broadly engaged across all risk-assessment behaviors, consistent with a generalized role in encoding anxiety-related information. In contrast, interneuronal subclasses exhibited striking functional specialization: parvalbumin (PV) interneurons were selectively recruited during uSAP and head dipping, behaviors associated with direct threat exposure, whereas somatostatin (Sst) interneurons were preferentially activated during pSAP, which reflect approach–avoidance conflicts and decision-making processes. Collectively, these findings establish a microcircuit-level framework in which distinct vH neuronal subclasses differentially gate risk-assessment strategies, enabling flexible transitions between avoidance and exploratory behaviors during anxiety.

## Introduction

Anxiety-related behaviors are fundamental adaptive responses that allow animals to evaluate potential threats and select context-appropriate actions. The ventral hippocampus (vH) has emerged as a central hub in anxiety, integrating spatial, sensory, and affective information to continuously monitor environments, evaluate potential threats, and shape ongoing approach–avoidance decisions through dynamic modulation of neural circuit function (*1–6*). The vH influences anxiety behavior through multiple functionally specialized pathways, including projections to the medial prefrontal cortex (mPFC) (*4, 7–9*), lateral septum (LS) (*10, 11*), and lateral hypothalamus (LH) (*5*), as well as through coordinated network dynamics and afferent control from the basolateral amygdala (*12, 13*). Moreover, vH projections to the nucleus accumbens encode motivational conflict and latent vulnerability states, shaping approach decisions under uncertainty and stress (*14–16*). These long-range projection pathways are not broadcasted uniformly but are selectively recruited and dynamically gated by local inhibitory microcircuits within the vH. Diverse interneuronal subclasses, including parvalbumin (PV) and somatostatin (Sst) interneurons, control pyramidal neuron activity through temporally precise perisomatic and dendritic inhibition, enabling behavior- and context-dependent routing of information. Recent work demonstrates that these inhibitory microcircuits differentially contribute to fear learning and anxiety, suggesting that local circuit dynamics and architecture within the vH underlie different forms of emotional behaviors (*1–3, 17, 18*). Together, these findings position the vH as a modular hub in which local microcircuits interact with brain-wide projection-specific pathways to regulate distinct components of adaptive behaviors.

Classic ethological studies, including the elevated plus maze (EPM), emphasize that anxiety is not a unitary construct, but instead comprises multiple, dissociable risk assessment and exploratory behavioral features (*19–21*). These include protected and unprotected stretch-attend posture (pSAP, uSAP) and head-dipping, as well as climbing, locomotion, and grooming, which reflect distinct ethologically defined anxiety-related behaviors (*22, 23*). Despite decades of behavioral work and clear pharmacological validity for many of these anxiety-related measures (*22, 24–26*), neuroscience experiments frequently reduce EPM behavior to coarse metrics (open arm time or entries), effectively treating anxiety as a scalar internal variable rather than a dynamic sequence of microstates or behavioral features (*27*). Consequently, how specific neural circuits and neuronal subclasses correspond to distinct explorative and risk-assessment behavioral features remains poorly understood. Bridging this gap between ethologically grounded behavioral features and circuit-level mechanisms is essential for a more mechanistic and biologically faithful understanding of anxiety.

In the present study, we combine high resolution 3D behavioral tracking and ethogram-based classification with cell-specific *in vivo* calcium imaging and temporally precise optogenetic manipulations to study vH microcircuits mechanisms of ethologically defined anxiety-related behaviors. Our results indicate that vH pyramidal neurons are widely recruited across all forms of risk-assessment behavior, supporting their broad role in representing anxiety-related information. By contrast, interneuron subclasses displayed marked functional specificity: PV interneurons were primarily active during uSAP and head-dipping, behaviors linked to immediate threat exposure, while Sst interneurons were preferentially engaged during pSAP, reflecting evaluation under approach–avoidance conflict and decision-making. Our results provide a novel framework for understanding how vH microcircuits orchestrate the spatiotemporal unfolding of anxiety-related features.

## Results

### Ethologically defined behaviors on the EPM

The EPM is a widely used assay for assessing anxiety-like behavior in rodents. It exploits the natural conflict between the drive to explore novel environments and the avoidance of open, elevated, and potentially threatening spaces. Mice display a complex repertoire of micro behaviors on the EPM that may reflect transient states as they traverse the environment. Standard 2D tracking of center-of-mass of mice on the EPM often fails to distinguish between distinct behavioral patterns, such as specific forms of exploration or threat evaluation. To resolve the fine microstructure of anxiety and systematically characterize the repertoire of anxiety-related behaviors on the EPM beyond coarse metrics (e.g., time in or entries to open arms), we implemented high-resolution 3D behavioral tracking combined with ethogram-based behavioral classification (*28*). During a 10-minute free-exploration session on the EPM, the mouse behavior was recorded at 30 Hz using three synchronized cameras, one overhead camera providing a complete top-view trajectory and two horizontal cameras aligned to the distal ends of the open arms to capture fine-scale postural dynamics (Fig. 1A). Six major ethologically defined behaviors were identified: grooming, walking, climbing, protected stretch-attend posture (pSAP), unprotected stretch-attend posture (uSAP) and head dipping (Fig. 1B,C; Movies S1-6). Grooming involves face and body cleaning and is generally considered an internally directed behavior. Walking reflects locomotion along the arms or through the center zone. Climbing consisted of rearing with the forepaws contacting the EPM wall and represents exploratory engagement with the environment. Head dipping involves the mouse extending and lowering its head over the edge of the open arm to inspect the environment below. Stretch-attend posture is characterized by a forward elongation of the body and tail, reflecting active environmental surveillance and heightened alertness. pSAP differs from uSAP in that during pSAP the mouse extends its body forward while keeping part of its body (typically the hindquarters) within the closed arm and center zone and quite often with part of the body touching the walls, whereas in uSAP the entire body is exposed onto the open arm(*22*).

**Fig. 1.**
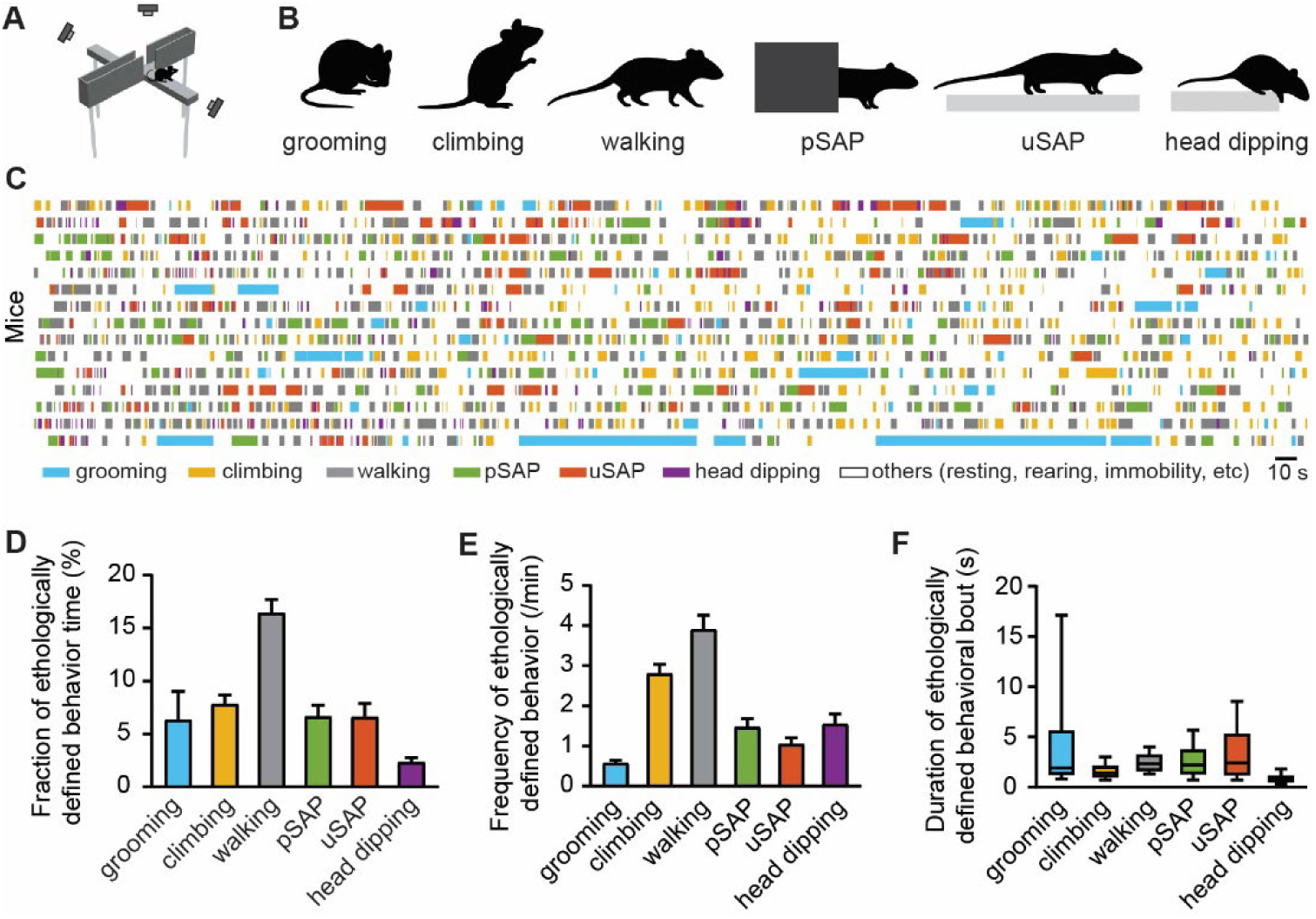
Ethologically defined behaviors on the EPM. (**A**) High-resolution 3D recording of mouse behaviors on the EPM. (**B**) Major ethologically defined behaviors identified on the EPM. (**C**) Ethograms during a 10-minute EPM test across 15 mice. (**D**) Time percentage of each ethologically defined behaviors during EPM testing. (**E**) Frequency of each ethologically defined behaviors per mouse. (**F**) Duration of ethologically defined behavioral bouts. n = 15 mice.

For each ethologically defined behaviors, we quantified their duration percentage of total EPM test time, occurrence frequency, and median bout duration. We observed that walking accounts for the greatest fraction of total EPM time (16.4%), whereas head dipping accounts for the least (2.3%). The other phenotypes including grooming, climbing, pSAP, and uSAP each lasted a comparable fraction of total EPM time (6-8%, Fig. 1D). Walking is also the most frequently occurring ethologically defined behavior, appearing on average 3.9 times per minute, followed by climbing at 2.8 times per minute (Fig. 1E). Grooming is the least frequent behavior, occurring only 0.6 times per minute. We next examined the median duration of behavioral bouts across mice and found that grooming showed substantial variability, whereas the durations of other ethologically defined behaviors were relatively stable (Fig. 1F). Notably, head dipping is the shortest ethologically defined behavior, with a median bout duration of 0.7 second, whereas the other phenotypes lasted around 2 seconds. This data indicate that mice express a stable and quantifiable set of distinct explorative and anxiety-related behaviors on the EPM, beyond simple open/closed arm occupancy metrics.

### Ethologically defined behaviors on the EPM are spatially and temporally structured

To capture how threat exposure modulates behavior, we first defined a spatial gradient across the EPM, ranging from the sheltered closed arms to the fully exposed open arms. Both the closed and open arms were equally divided into four squared 8cm × 8cm sub-zones, yielding a total of 17 zones when added to the center zone. In the closed arm, zones from C1 to C4 reflected increasing levels of safety, with C4 at the distal end surrounded by three walls representing the safest region. C1, although still part of the closed arm and therefore a rather safe zone, positions the mouse at the boundary where the external open space becomes fully visible, often triggering exploratory decisions toward the more anxiogenic open arms. The open arms were similarly divided into four sub-zones (O1 - O4) based on exposure and distance from the closed arm. O4 represents the most exposed and thus dangerous region and requires the longest retreat path to safety zones. Together, this segmentation defined a spatial gradient incorporating distinct safety and threat components, from the closed to the open arms of the EPM (Fig. 2A).

To test whether ethologically defined behaviors were uniformly distributed across the EPM, we computed the total time spent and the count of behavioral events per ethologically defined behaviors in each zone (Fig. 2B). We found that grooming and climbing occurred predominantly in C4, indicating self-maintenance and exploratory behavior in safe zones. Walking occurred predominantly in the closed arms, either when the mouse traveled within a single closed arm or transitioned from one closed arm to another through the center zone. pSAP were significantly enriched at the boundaries between the closed and open arms (zones C1, CE, and O1), reflecting active environmental surveillance, anticipation of potential threat, risk assessment and decision-making processes regarding whether to approach to or avoid the anxiogenic open arms. Interestingly, uSAP occurred for longer durations in O4 but with higher frequency in O1, suggesting that mice perform uSAP more often, but briefly, in less anxiogenic regions, whereas in highly anxiogenic zones they exhibit uSAP less frequently, but once initiated, the behavior persists longer. Head dipping occurred across all open-arm zones (O1 to O4) but was most frequent in O1. In contrast to pSAP, which support risk assessment within a protected context, uSAP and head dipping represent risk assessment that occurs through direct exposure to potential threat while anxiety remains elevated.

Next, we applied a Markov chain analysis across the six ethologically defined behaviors on the EPM to examine their temporal sequence. For each behavior, we annotated the initiation and termination time points of every bout. A custom algorithm was then used to temporally order all ethologically defined behavioral bouts for each mouse, after which we computed the number of transitions from one ethologically defined behavior to another (Fig. 2C). For example, walking can not only transition to other ethologically defined behaviors, but also recur as walking itself. The transition probabilities among ethologically defined behaviors vary widely, reflecting the underlying correlation and functional relationships between distinct behavioral microstates. Each ethologically defined behavior has a unique transition profile, indicating how likely it is to lead to exploratory or anxiety-related behaviors. Importantly, the outgoing transition probabilities from any given ethologically defined behavior sum to 100%, providing a normalized measure of how behavioral sequences are organized and how mice dynamically shift between behavioral microstates during the EPM test (Fig. S1A). The Markov chain analysis revealed extensive and complex transitions among all ethologically defined behaviors (Fig. S1B). To reveal the major transition trends among ethologically defined behaviors, we applied a threshold mask that filtered out probabilities below 25%, retaining only high-probability transitions (Fig. 2D). Overall, walking emerged as a central hub connecting all other ethologically defined behaviors. After completing any given behavior, mice typically engaged in brief walking before initiating the next ethologically defined behavior. This analysis identified three distinct transition clusters: cluster 1 comprising climbing, grooming, and walking; cluster 2 comprising uSAP and head dipping; and cluster 3 consisting solely of pSAP. This analysis demonstrated stereotyped sequences of ethologically defined behaviors rather than random switches.

Grooming predominantly transitioned to either walking or climbing. Walking and climbing also showed high recurrence and strong bidirectional transitions. Together, these three ethologically defined behaviors form an exploratory behavioral cluster, expressed primarily while mice remain within the safe closed arms. uSAP and head dipping also exhibited substantial bidirectional transitions. Notably, 48% of uSAP events were followed by a head dip, indicating that after evaluating the openness of the environment, mice often proceed to assess height-related threat. Head dipping showed a 32% recurrence probability, consistent with rapid, repeated dipping that facilitates thorough threat assessment. Because both uSAP and head dipping occur mainly on the open arms, this second cluster reflects ethologically defined behaviors associated with risk engagement or anxiety processing when the mice are fully exposed to an anxiogenic environment. Strikingly, pSAP formed a relatively isolated cluster, with its strongest outbound transition directed toward walking (46.3%). As pSAP occurs primarily in the EPM decision-making zone and at safety/danger boundaries (C1, CE, and O1), we interpret this risk assessment behavior as reflecting potential threat prediction and approach-avoidance related decision-making, wherein the mouse must choose either to retreat to the safe closed arms or to proceed toward the anxiogenic open arms, requiring a transition probability to walking.

**Fig. 2.**
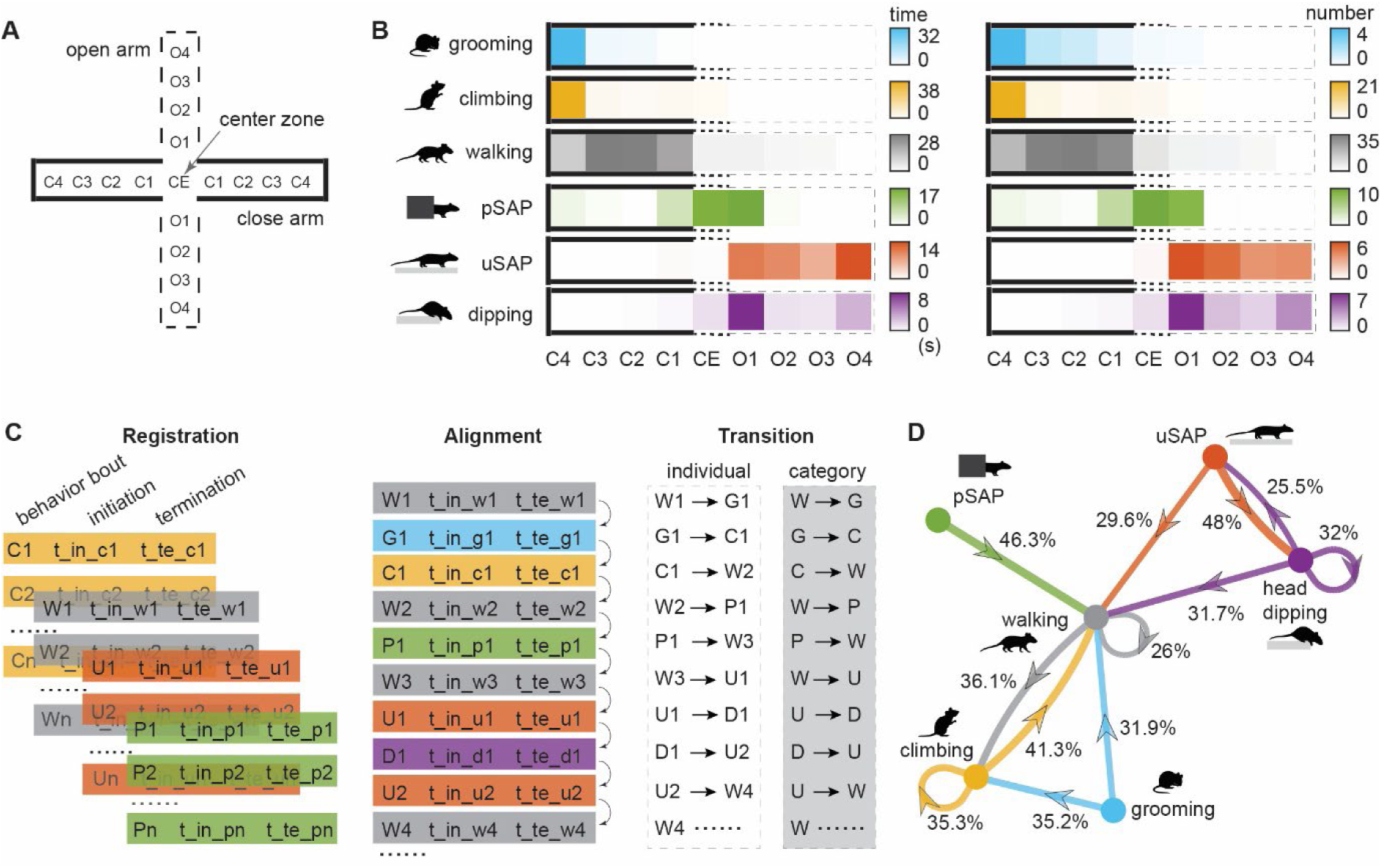
Ethologically defined behaviors on the EPM are spatially and temporally structured. (**A**) The EPM was segmented into 17 zones capturing a safety to risk gradient from the closed arms to the open arms. (**B**) Time spent and number of ethologically defined behaviors occurring in each EPM zone. (**C**) Workflow of Markov chain analysis among ethologically defined behaviors. (**D**) High-probability transitions among six major ethologically defined behaviors on the EPM. Transition probability from one behavioral microstate to another is shown in percentage. Arrows indicating the transition direction from one ethologically defined behavior to another. n = 15 mice.

Taken together, our analyses demonstrate that ethologically defined exploratory and anxiety-related behavioral are not expressed randomly, but are systematically organized across spatial and temporal dimensions. Spatially, they align along the EPM’s safety to risk gradient, with self-maintenance and exploratory microstates such as grooming, walking and climbing confined to closed arms; risk assessment and decision-making related to pSAP concentrated around center zones; and anxiety-related but risk-engaging uSAP and head dipping emerging on the open arms. Temporally, they unfold in structured and stereotyped transition sequences, revealing functionally distinct behavioral clusters and directional flow among ethologically defined.

### Distinct subclasses of vH neurons are activated by ethologically defined behaviors

What are the neuronal substrates and circuit basis underlying these ethologically defined behaviors? The vH has been implicated in anxiety-related behaviors (*29*). Accordingly, we targeted the CA1 subfield of the vH to monitor activity from identified neuronal subclasses using a GRIN lens and a head-mounted miniature microscope in freely behaving mice (Fig. 3A, S2A). The genetically encoded Ca²⁺ indicator GCaMP6f was expressed in vH pyramidal neurons (Fig. 3B) and, in separate experiments, in local PV and Sst interneurons using Cre-dependent AAV virus-mediated expression in PV.Cre or Sst.Cre mouse lines (Fig. 3C,D). Calcium traces from individual neurons were aligned to ethogram-defined behavioral bouts to relate single-cell activity to specific ethologically defined behaviors.

Pyramidal neurons constitute the principal excitatory population, integrating convergent synaptic inputs and serving as the primary source of long-range hippocampal output to distributed brain regions. We first examined how vH pyramidal neurons participate to the six ethologically defined behaviors (Fig. 3E). Pyramidal neurons were minimally activated during walking (4.5%). Although locomotion is a strong driver of activity in the dorsal hippocampus, particularly in place cells, vH pyramidal neuron population activity remained low during walking. This is consistent with the vH’s specialization in encoding stimulus features and internal behavioral states rather than locomotor output (*30, 31*). A small subset of pyramidal neurons (10%) was recruited during grooming, likely reflecting vH’s role in representing internal state transitions and perceived safety. Climbing elicited 15.5% recruitment, which is part of the broader exploratory repertoire and may correspond to increased vigilance and sensory information sampling (*32*). Notably, pyramidal neurons showed markedly higher activation during ethologically defined anxiety-relevant behaviors, including pSAP, uSAP, and head dipping (15%, 19%, and 26.5%, respectively). The progressively larger fractions of activated neurons mirror the increasing anxiety intensity ‘load’ and threat exposure associated with these behaviors, highlighting the role of vH pyramidal neurons in representing approach-avoidance conflict and anxiogenic states (*6*).

To understand how pyramidal neuron activity patterns are controlled at the microcircuit level, we next focused on interneurons (*33*). Among the major GABAergic inhibitory subclasses, PV interneurons are of particular interest because they exert powerful perisomatic inhibition and precisely regulate the timing and gain of pyramidal neuron firing (*34–37*). Recent studies suggest that PV interneurons modulate pyramidal neurons activity during anxiety behavior (*1, 2, 38*). However, these findings do not reveal how PV interneurons engage during specific ethological behaviors that make up an integrated anxiety response (e.g., SAP, head dipping). Yet, it remains unknown whether PV interneurons exhibit specific recruitment pattern during anxiety-related ethological behaviors or whether their modulation is uniform across different components of the anxiety behavior repertoire. To address this gap, we used the same *in vivo* calcium imaging method employed for pyramidal neurons and monitored PV interneuron activity aligned to ethologically defined behavioral bouts. This allowed us to determine whether PV interneurons exhibit selective responses to discrete ethologically defined behaviors or instead show global, state-dependent modulation across exploration and avoidance.

Our results show that vH PV interneurons are recruited to a similar extent as pyramidal neurons during grooming, climbing, and walking, suggesting a role in general network stabilization and modulation of pyramidal neuron activity during routine and explorative behaviors. However, unlike pyramidal neurons, PV interneurons were minimally active during pSAP, but were strongly recruited during uSAP and head dipping (Fig. 3F). This pattern indicates that PV interneurons are preferentially involved in ethologically defined behaviors associated with immediate threat exposure and risk engagement. In contrast, PV activity is less prominent during pSAP, suggesting a minor function in risk assessment and decision-making processes. Together, these findings suggest that vH PV interneurons selectively shape pyramidal neuron output in response to proximal threat without directly influencing decision-making in the transition between the safe and anxiogenic compartment of the EPM.

Whereas PV interneurons predominantly control perisomatic excitability, Sst interneurons target pyramidal neuron dendrites to regulate synaptic integration and input-output gain (*39–45*), potentially exerting behavioral state-dependent control over pyramidal neuron activity. Sst interneurons showed modest recruitment during grooming, walking, and climbing, similar to the small subsets of pyramidal and PV neurons engaged during these behaviors. However, Sst interneurons exhibited markedly increased activation during SAP behaviors. A larger fraction of Sst cells was active during pSAP (28%) than uSAP (21%), indicating stronger Sst recruitment in risk assessment and approach-avoidance related decision-making. During head dipping, Sst population activation level was low, with only 9.3% of neurons activated (Fig. 3G).

**Fig. 3.**
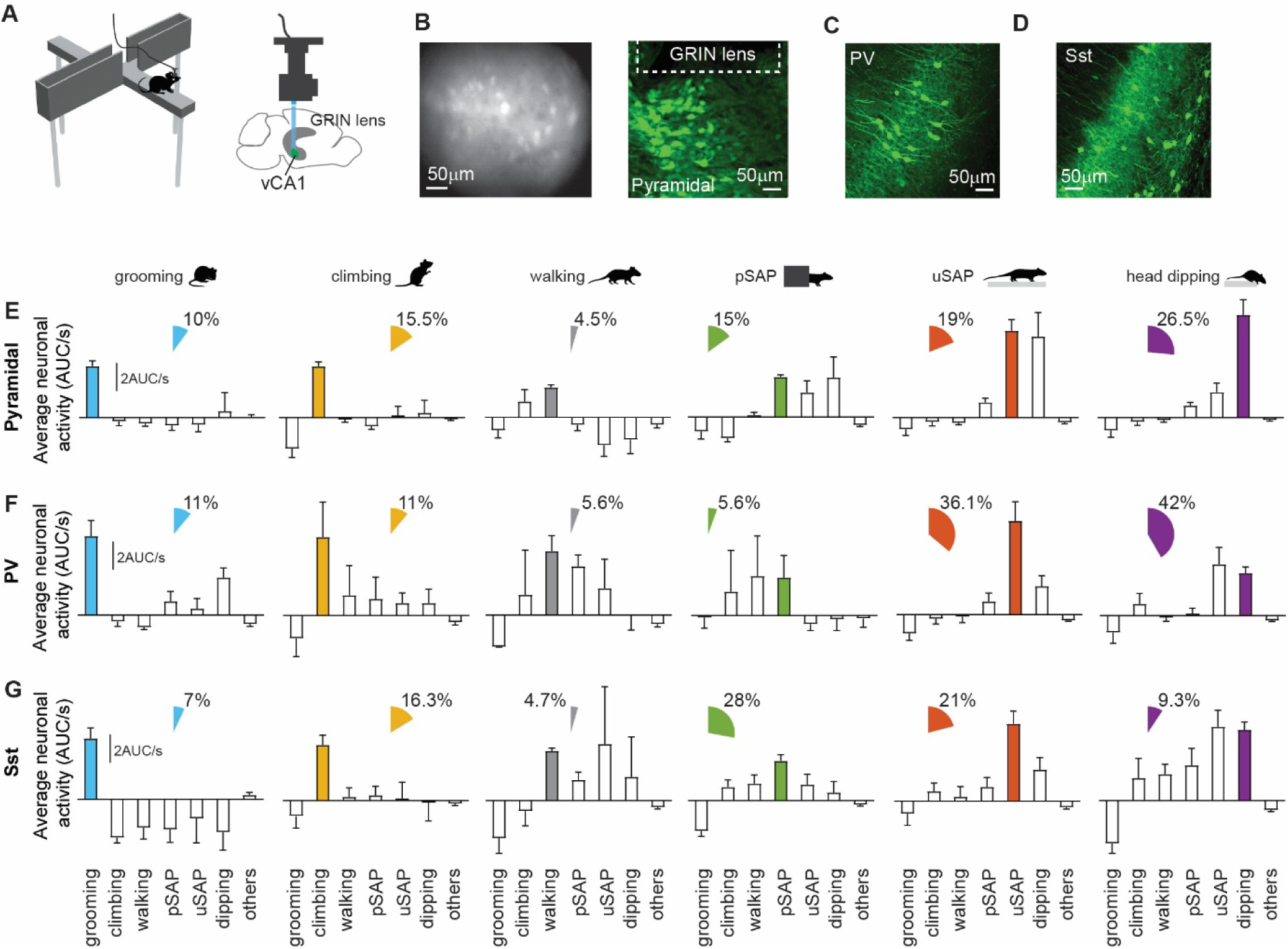
Distinct subclasses of vH neurons are activated by ethologically defined behaviors. (**A**) Schematic showing *in vivo* calcium imaging in vH of freely moving mice during EPM exploration by using a miniscope. (**B**) Representative images of GCaMP6f expressing pyramidal neurons acquired by miniscope *in vivo* (left) and confocal imaging *in situ* (right). (**C**,**D**) Representative confocal images of PV and Sst interneurons in vH. (**E-G**) Average neuronal activity and the fraction of neurons selectively recruited during ethologically defined behaviors. (E) n = 200 pyramidal neurons from 5 C57BL/6 mice; (F) n = 36 neurons from 4 PV.Cre mice; (G) n = 43 neurons from 5 Sst.Cre mice.

In a complementary anxiety test, the forced anxiety shifting test (FAST), we observed a similar recruitment pattern of inhibitory interneuron subclasses. PV interneurons were robustly and preferentially recruited during uSAP and head-dipping behaviors, whereas Sst interneurons showed comparatively weaker engagement (Fig. S3). Notably, the FAST involves rapid transitions between closed and open environments (*1*), which effectively preclude the emergence of the pSAP phenotype. This constrained geometry biases behavior toward immediate exposure to height and bright openness. Under these conditions, Sst interneuron recruitment during head dipping was modestly elevated relative to EPM, potentially reflecting the higher and more sustained anxiogenic load imposed by this forced exposure paradigm. Nevertheless, across both FAST and EPM, PV interneurons consistently exhibited stronger recruitment than Sst interneurons during uSAP and head dipping. This cross-task consistency supports the conclusion that PV interneurons are preferentially engaged during uSAP and head dipping behavioral microstates associated with immediate environmental exposure to threat, rather than task-specific features of the behavioral apparatus.

Altogether, we observed distinct activation profiles associated with specific ethologically defined behaviors across pyramidal neurons, PV and Sst interneurons of the vH. Pyramidal neurons showed progressively higher activation from grooming, climbing and walking, to pSAP, uSAP and head dipping behavioral phenotypes. PV interneurons showed strong activation during uSAP and head dipping, while Sst interneurons displayed the highest recruitment during pSAP, followed by uSAP, with limited engagement during head dipping.

### Subclasses of vH neurons are differentially activated at the initiation of ethologically defined anxiety-related behaviors

After defining overall recruitment patterns of vH neuronal subclasses, we examined how these neurons are engaged as ethologically defined anxiety-related behaviors emerge, since time-locked neural activity may reflect circuit mechanisms that precede and potentially drive these behaviors. We aligned calcium transients to the initiation of each behavioral bout and extracted fluorescence traces from -1.5 s to +1.5 s windows around the onset of ethologically defined anxiety-related behaviors. Z-score normalized traces were used to compute pre- and post-onset area-under-the-curve (AUC) values for pSAP, uSAP, and head dipping.

pSAP-, uSAP- and head dipping-selective pyramidal neurons (Fig. 3E) exhibited a significant increase in activity after behavioral initiation (Fig. 4A-C). This consistent, time-locked recruitment across all three behaviors suggests that vH pyramidal neurons generally encode anxiety-related microstates ranging from risk assessment in safer to more anxiogenic locations. Pyramidal neurons probably provide a feed-forward drive to downstream brain structures during the initiation and maintenance of ethologically defined anxiety-related behaviors. PV interneurons displayed a rapid rise in activity after the onset of uSAP and head dipping (Fig. 4E,F), whereas very few PV interneurons showed high but largely variable activity during pSAP (Fig. 3F, 4D). This is consistent with their fast perisomatic inhibition and known role in modulating pyramidal neuron output in anxiogenic environment (*1, 2*). These data indicate that PV interneurons contribute minimally to risk assessment and decision making in safer zones, but strongly to ethologically defined anxiety-related behaviors involving direct threat exposure in anxiety-inducing zones of the EPM open arms. In contrast, Sst interneurons showed a clear activation peak at the onset of pSAP, but not at the onset of uSAP or head dipping (Fig. 4G-I). Although a substantial fraction of Sst neurons was active throughout the duration of uSAP, their activation was more sustained and slowly evolving, rather than time-locked to the initiation of behavior (Fig. 3G, 4H).

**Fig. 4.**
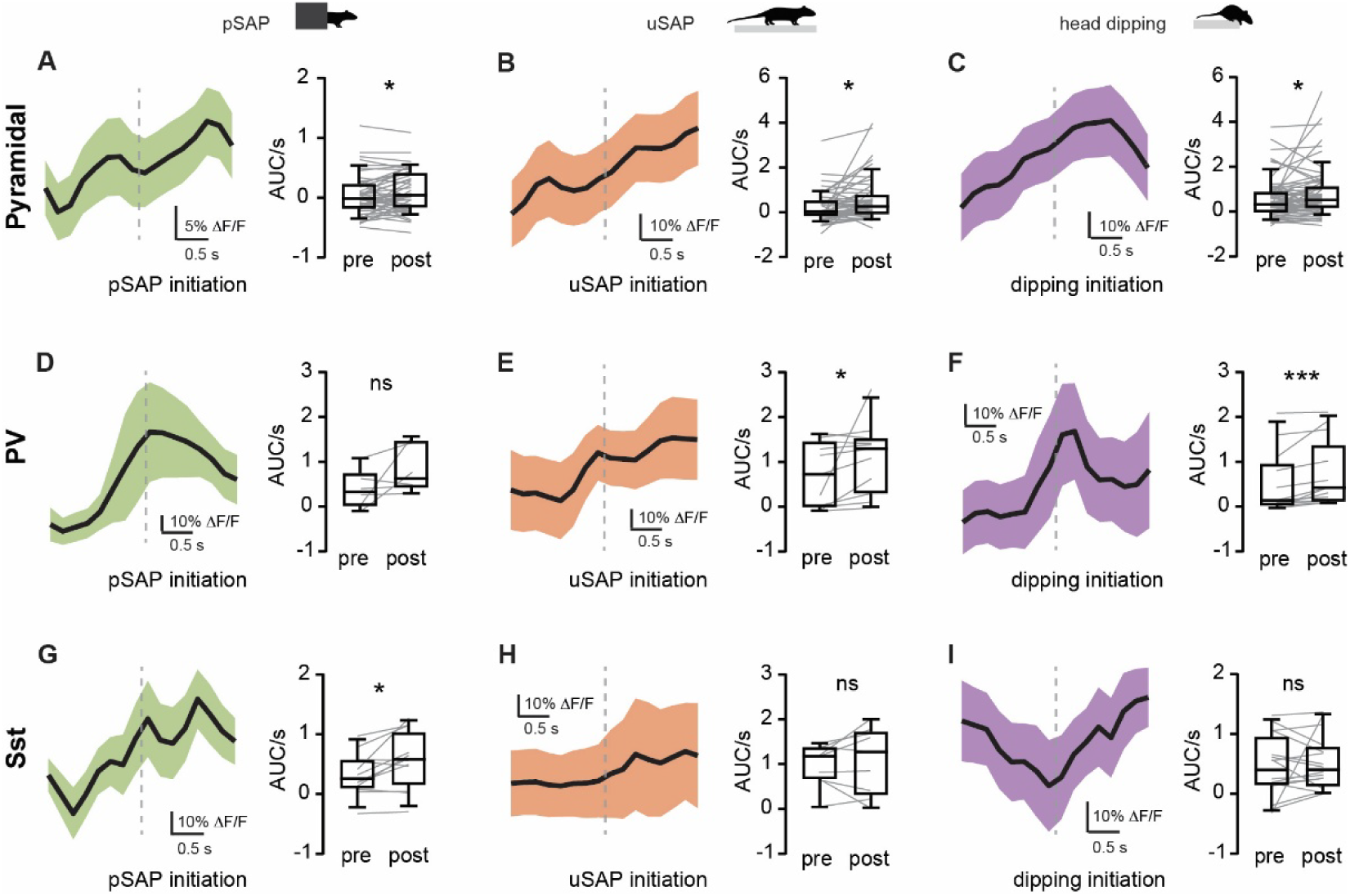
Subclasses of vH neurons are differentially activated at the initiation of ethologically defined anxiety-related behaviors. Averaged normalized activity at the initiation of pSAP (**A,D,G**), uSAP (**B,E,H**) and head dipping (**C,F,I**) with recruited pyramidal neurons (**A,B,C**), PV interneurons (**D,E,F**) and Sst interneurons (**G,H,I**). Box plots showing 1.5 seconds AUC before and after behavior initiation. A, n=51 cells, 5 mice; B, n=54 cells, 5 mice; C, n=66 cells, 5 mice; D, n=6 cells, 4 mice; E, n=11 cells, 4 mice; F, n=11 cells, 4 mice; G, n=12 cells, 4 mice; H, n=9 cells, 4 mice; I, n=16 cells, 4 mice. Paired t-test, *p < 0.05, ***p < 0.001. ns, not significant.

### Optogenetic inhibition of vH neuronal subclasses differentially affects ethologically defined anxiety-related behaviors

To determine whether these neuronal subclasses are necessary for specific ethologically defined anxiety-related behaviors, we performed cell population specific optogenetic inhibition during EPM center and open arm exploration using the eNpHR3.0 opsin. Silencing pyramidal neurons significantly increased the duration and frequency of uSAP and head dipping, without affecting pSAP (Fig. 5A-C, S2B). Inhibiting PV interneurons reduced uSAP and head dipping (Fig. 5D-F), an opposite pattern compared to pyramidal neuron inhibition. By using a dual-channel miniscope (*1, 46*), we observed elevated pyramidal neuron activity during optogenetic inhibition of PV interneurons (Fig. 5J,K, S2C). These data demonstrate a functional vH microcircuit involving pyramidal neurons and PV interneurons that governs uSAP and head dipping behaviors.

Silencing Sst interneurons specifically reduced both the duration and frequency of pSAP, while leaving uSAP and head dipping unchanged (Fig. 5G-I, S2B). These results align with the neuronal activity data (Fig. 4G) and indicate that Sst interneurons and pyramidal neurons (Fig. 4A, 5L, S2C) promote risk assessment and decision-making behaviors, particularly in transitional zones where animals evaluate potential threats while staying in a safer environment.

**Fig. 5.**
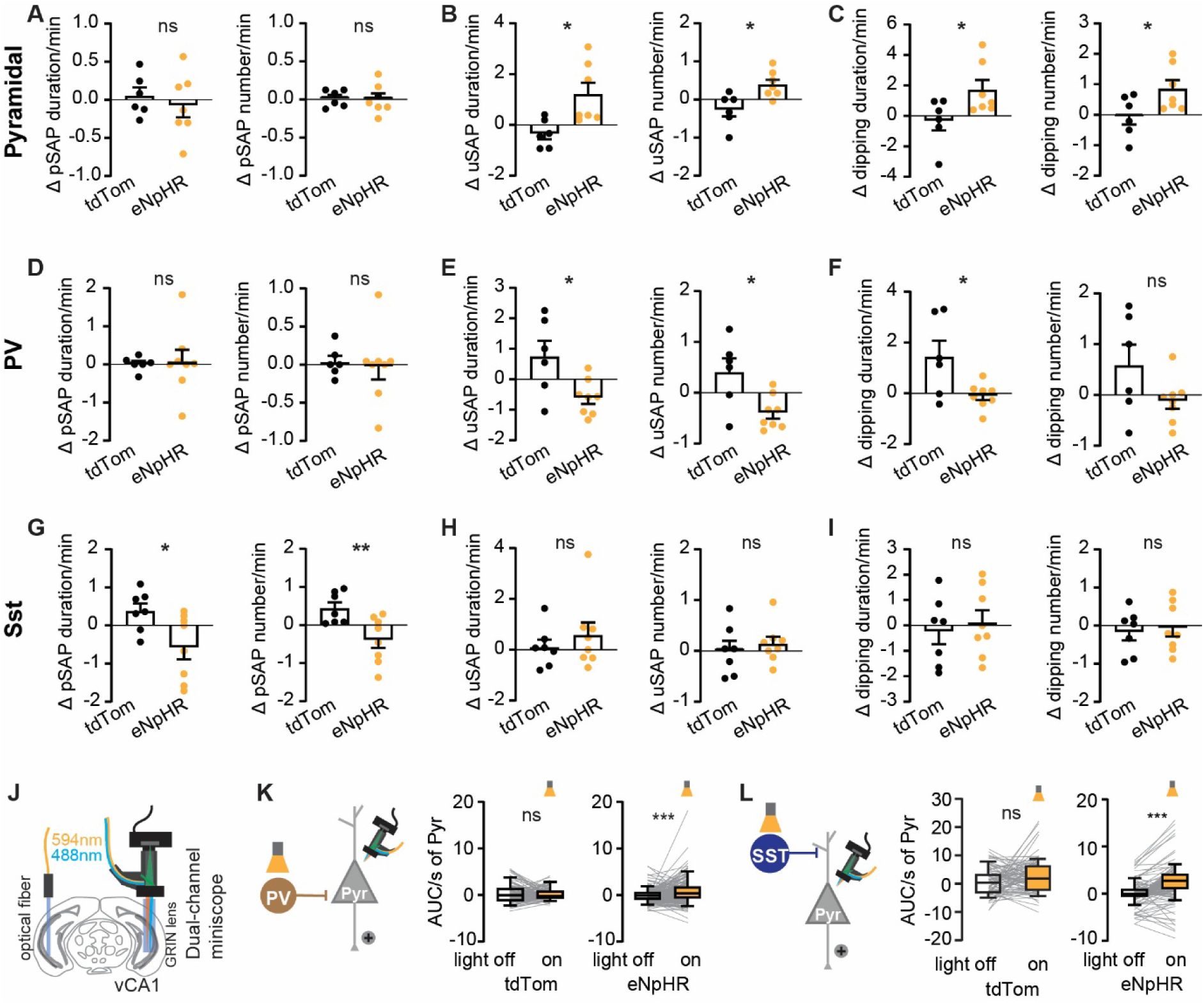
Optogenetic inhibition of vH neuronal subclasses differentially affects ethologically defined anxiety-related behaviors. (**A-C**) Optogenetic inhibition of pyramidal neurons had no effect on pSAP (A) and increased duration and frequency of uSAP (B) and head dipping (C). tdTomato n = 6 mice; eNpHR3.0 n = 7 mice. (**D-F**) Optogenetic inhibition of PV interneurons reduced duration and frequency of pSAP (D) and had no impact on uSAP (E) and head dipping (F). tdTomato n = 7 mice; eNpHR3.0 n = 8 mice. (**G-I**) Optogenetic inhibition of Sst interneurons had no effect on pSAP (G) and reduced duration and frequency of uSAP (H) and head dipping (I). tdTomato n = 6 mice; eNpHR3.0 n = 8 mice. (**J**) Design of dual-channel miniscope enabling bilateral optogenetic manipulation and calcium imaging. (**K**) Optogenetic inhibition of PV interneurons disinhibit pyramidal neuron activity. tdTomato n = 89 neurons from 2 mice; eNpHR3.0 n = 125 neurons from 4 mice. (**L**) Optogenetic inhibition of Sst interneurons disinhibit pyramidal neuron activity. tdTomato n = 82 neurons from 4 mice; eNpHR3.0 n = 130 neurons from 5 mice. (A-I) Unpaired t test and (K,L) Wilcoxon matched-pairs signed rank test, *p < 0.05, **p < 0.01, ***p < 0.001. ns, not significant.

Together, the onset-aligned calcium transients during ethologically defined anxiety-related behaviors and the optogenetic inhibition experiments indicate that distinct vH neuronal subclasses exert causal and dissociable influences on anxiety-relevant behaviors. These results reveal that vH pyramidal neurons, PV interneurons, and Sst interneurons differentially orchestrate anxiety microstates as a function of changes in threat exposure.

## Discussion

In animal models, anxiety is often operationalized as a stable internal state inferred from spatial avoidance on the EPM. However, anxiety can also be understood as a dynamic process that continuously regulates whether exploration is initiated, paused, or terminated in response to uncertainty and threat. In this study, we decompose anxiety-related behavior on the EPM into discrete, ethologically grounded behavioral microstates and map these onto the dynamics of defined vH neuronal subclasses. By capturing ongoing behavioral sequences and transitions rather than the aggregate open arm occupancy and entry, this approach reframes anxiety as a dynamic process unfolding through structured behavioral microstates. We show that the vH does not regulate anxiety as a unitary internal state. Instead, pyramidal neurons, PV, and Sst interneurons are recruited in parallel to control distinct components of risk assessment and engagement, decision-making and anxiety regulation. These findings support a distributed microcircuit architecture in which PV- and Sst-based inhibitory microcircuits differentially shape specific ethologically defined anxiety-related behaviors rather than globally modulating anxiety levels.

### Anxiety as a structured sequence of behavioral microstates rather than a unitary state

Traditional EPM analyses collapse behavior across time and space, implicitly assuming that anxiety can be captured by aggregate measures such as open arm occupancy and entries. Our results challenge this assumption by demonstrating that mice express multiple, stereotyped ethologically defined behaviors, such as grooming, walking, climbing, pSAP, uSAP, and head dipping that differ in their spatial context, temporal structure, and associated neuronal activity.

These ethologically defined behaviors are not interchangeable proxies of a single anxiety dimension but reflect qualitatively distinct modes of interaction with the environment.

Crucially, many anxiety-relevant behaviors occur within zones that are conventionally categorized as safe, particularly near boundaries such as C1 zone on the EPM defined in this study where animals must decide whether to increase exposure. This observation underscores that anxiety is also salient at decision boundaries where the animal weighs potential threat while physically still mostly in the safe environment. By explicitly modeling ethologically defined behaviors along a graded safety to risk axis, our framework reveals how anxiety emerges through transitions between behavioral states rather than through sustained occupancy of an anxiogenic space.

This framework also enables direct alignment between specific behavioral microstates and neuronal activity, demonstrating how vH circuits are differentially engaged during protected risk assessment, exploration, and unprotected threat exposure. Importantly, this structure also exposes rich transitions between ethologically defined behaviors. For example, bidirectional transitions between uSAP and head dipping, as well as structured relationships among grooming, climbing, and walking, are largely overlooked by traditional analyses yet likely reflect core features of anxiety to resolve approach-avoidance conflict over time. Although not all observed transitions were analyzed in depth here, their systematic coupling to neuronal dynamics suggests that anxiety emerges and vanishes through structured state changes implemented by local microcircuits. This framework therefore provides a generalizable approach for studying anxiety as a temporally organized risk assessment and decision-making processes, rather than a static emotional state.

### Functional dissociation among vH microcircuits in ethologically defined anxiety-related behaviors

At the circuit level, our data indicate that the vH does not exert uniform control over anxiety-relevant behaviors. Instead, pyramidal neurons, PV and Sst interneurons are differentially engaged across ethologically defined anxiety-related behaviors and exert distinct causal influences. Pyramidal neurons and PV interneurons were preferentially recruited during behaviors involving direct risk engagement, including uSAP and head dipping. In contrast, Sst interneurons showed selective recruitment during pSAP and were causally required for its maintenance. Sst activity appears necessary to stabilize a behavioral state in which exploration is constrained while contextual information is gathered to control the decision making process of entering or not into anxiogenic zones.

The selective dependence of pSAP on Sst interneurons suggests that risk assessment is not a passive intermediate between movement states but an actively maintained anxiety-related process. pSAP occurs in emotional conflict zones on the EPM (i.e., including C1, CE and O1) in which approach and avoidance drives are engaged. The fact that optogenetic Sst inhibition reduced pSAP implies that Sst interneurons are required to sustain the spatial and temporal structure of risk assessment in decision making zones. Within the vH, Sst interneurons may implement this function by regulating dendritic integration in pyramidal neurons, thereby increasing the threshold for behavioral switching and promoting extended contextual information gathering. This interpretation is consistent with the relatively weak recruitment of PV interneurons during pSAP, suggesting that PV-mediated perisomatic inhibition is not a core components of risk assessment in decision making zones and becomes relevant primarily in zones with direct threat exposure such as during uSAP and head dipping. This functional dissociation supports a model in which ethologically defined anxiety-related behavior arises from the interaction of distinct inhibitory circuit mechanisms, rather than from a single unified one.

### Caveats in linking ventral hippocampal activity to anxiety-related behavior

Optogenetic manipulations reveal a selective requirement for ventral hippocampal (vH) pyramidal neuron activity in uSAP and head dipping, but not in pSAP. Although a subset of pyramidal neurons showed elevated activity during pSAP (Fig. 4A), inhibiting pyramidal output did not affect either the duration or frequency of pSAP bouts (Fig. 5A). This dissociation indicates that pyramidal neuron activity during pSAP is not necessary for maintaining this behavioral state. Instead, pyramidal engagement may reflect state representation or coordination, potentially broadcasting contextual or threat-related information to downstream regions such as the prefrontal cortex or amygdala, rather than exerting local control over behavioral execution.

In contrast, local vH inhibitory circuitry appears to play a central role in stabilizing pSAP. The persistence of pSAP following pyramidal inhibition suggests that once risk assessment is engaged, its maintenance depends primarily on inhibitory mechanisms rather than on pyramidal output. We therefore propose a division of labor in which pyramidal neurons encode or signal the current behavioral state, while inhibitory interneurons determine whether that state persists or transitions.

A related pattern emerged from analysis of PV interneurons. While a small population of PV cells showed a nonsignificant trend toward increased activity during pSAP (Fig. 4D), optogenetic inhibition of PV interneurons did not alter pSAP expression (Fig. 5D), further supporting the idea that risk assessment is not governed by fast perisomatic inhibition. However, PV inhibition significantly reduced the duration, but not the frequency, of head dipping bouts (Fig. 5F). This selective effect suggests that PV interneurons regulate the sustainment of head dipping once initiated, implicating PV-mediated inhibition in maintaining this behavioral state rather than triggering its onset.

Together, these findings reveal a temporal and state-dependent uncoupling between neuronal activation and behavioral necessity. Nevertheless, a limitation of the current optogenetic approach is that spatially defined inhibition may miss brief windows of causal influence occurring at behavioral transitions. Closed-loop strategies (*47–49*), in which manipulations are triggered by the onset or offset of defined behaviors, will be critical to determine whether pyramidal and PV neurons causally regulate anxiety by shaping state transitions rather than sustained behavioral expression.

### Translational implications and microcircuit-targeted intervention

The ethologically defined anxiety-related behaviors identified in this study provide a mechanistic framework for understanding pathological anxiety in humans (*50, 51*). From a translational perspective, our findings suggest a shift away from globally acting anxiolytics which are often associated with multiple undesired effects (*52*) toward interventions that selectively target specific circuit motifs with distinct anxiety-related functions. The pSAP anxiety-related behavior closely mirrors human anxious indecision or hypervigilance, where individuals continuously monitor their environment and gather information while staying within a perceived safe space. The selective recruitment of Sst interneurons in pSAP may suggest an important role of vH Sst microcircuit in pathological indecision and persistent risk assessment and highlights a potential therapeutic target that is more specific than the broad inhibitory actions of classical anxiolytics such as benzodiazepines. Conversely, the strong engagement of PV interneurons and pyramidal neurons during uSAP and head dipping points to a complementary axis with particular relevance for anxiety disorders dominated by cue-and context-specific avoidance, such as specific phobias. Tools for tuning PV interneuron mediated inhibition, either pharmacologically or through circuit-level interventions, could therefore ‘renormalize’ approach/avoidance threshold in the presence of phobic stimuli without compromising the Sst-dependent capacity for cautious, adaptive risk assessment that remains essential in genuinely threatening contexts.

More broadly, the methodological framework introduced here enables anxiety to be quantified in terms of behavioral state occupancy, transition probabilities, and decision threshold which are more directly comparable parameters across species and potentially more relevant to human psychopathology than traditional avoidance metrics.

In summary, this study establishes a circuit-level framework in which anxiety emerges from dynamic interactions among behavioral microstates regulated by distinct vH microcircuits. By decomposing behavior into ethologically grounded phenotypes and linking these to causal neuronal mechanisms, we show that the vH functions not as an anxiety ‘ON/OFF’ switch but as a rather sophisticated detector of uncertainty and threat proximity. Future work combining projection-specific recordings and closed-loop perturbations will be essential for determining how these local computations are transformed into adaptive or maladaptive action by vH downstream networks. Finally, by resolving anxiety into microcircuit regulated behavioral phenotypes, our findings offer a translational framework for linking defined hippocampal circuit dynamics to distinct human anxiety symptoms, including maladaptive risk assessment, persistent indecision, and exaggerated avoidance, and suggest that interventions targeting these microcircuits may achieve symptom-specific modulation beyond global anxiolysis.

## Materials and Methods

### Animals

Mice aged 3 to 6 months were group housed with ad libitum access to food and water on a 12-hour light/dark cycle at constant temperature (22 ± 1 °C) and humidity (30 - 40%). C57BL/6J mice (Janvier Labs, France), PV.IRES.Cre (Jackson Laboratory, Strain #008069), Sst-IRES-Cre (Jackson Laboratory, Strain #013044). These transgenic mouse lines are of B6 background and had been backcrossed with C57BL/6J wild type mice for more than ten generations to produce heterozygous offspring with B6J congenic background. Only heterozygous mice were used for experiments. Following surgery, mice undergoing *in vivo* Ca^2+^ imaging experiments were single-housed to secure miniscope implantation. For optogenetic manipulation and rabies tracing experiments, littermates were randomly assigned to experimental conditions and group-housed with two to four mice per cage. All experimental procedures were performed in accordance with the guidelines of the Animal Welfare Office at the University of Bern and approved by the Veterinary Office of the Canton of Bern.

### Behavior

The mice were habituated to the room for at least 3 days before behavior test by placing mice in home cage into the testing chamber for 2 × 30 min daily. Each mouse had its own custom-made nesting house made of hard paper, which was used for transferring mice to testing apparatus to reduce mouse grabbing. The mice were also habituated to be moved out of the home cage many times prior to experiments.

### Elevated plus maze (EPM)

The EPM was custom-made and consisted of four arms. Each arm is 8 cm wide and 35 cm long and elevated 75 cm above the floor. The two opposing closed arms were enclosed by 18 cm high walls, whereas the other two were left open. The center zone is in the middle with a size of 8 cm × 8 cm. Prior experiments, mice were introduced into the middle part of a closed arm with head positioning toward the center zone. Then mice explored the EPM freely for 10 minutes in a sound attenuating chamber under 300 lux from roof-top light. Three high-resolution cameras operating at 30 Hz were used to record animal behavior. One camera was positioned above the EPM to capture movements from a top view, while two additional cameras were placed 15 cm away from the maze, aligned at the same height as the two open arms, to capture side views. All cameras were synchronized using OBS Studio to ensure precise temporal alignment across recordings. Animal positions were tracked using ANYmaze (Stoelting, USA). The mouse body center was used to define its location in closed arm, center zone or open arm of the EPM.

### Forced Anxiety Shifting Task (FAST)

The FAST paradigm is a trial-based anxiety behavioral task consisting of an elevated platform (160 cm from the ground, 10 cm×10 cm) surrounded by a motorized black cubicle (11 cm×11 cm). Mice were placed on top of the platform inside the black cubicle. The cubicle surrounding the platform was controlled by a linear motor (LinMot NTI AG) with a custom-made (Electronic Workshop, University Bern) digital input/output interface, which triggered a TTL to control the cubicle moving up and down. The FAST enabled us to submit mice to sequential and separate time-locked exposures to an anxiogenic situation (an elevated, bright, and open space), a novel situation (a dark green cubicle) or a safe situation (a closed and darker cubicle).

The FAST behavioral test consists of 5 ‘novelty control’ trials and 10 ‘anxiety test’ trials. To avoid photon bleaching during interneurons imaging, we decided to minimize novelty controls to 5 trials to ensure that good quality of data can be obtained in the anxiety test trials. In novelty control trials, mice were first held for 20 s in the closed compartment with black background walls (safe context). Subsequently, the black cubicle was lowered, exposing mice to the green cubicle (novel context) for 30 s. During anxiety test trials, mice were held for 20 s in the closed compartment (safe). Afterwards, the black cubicle slid down to the platform base level, forcibly exposing mice to the high and brightly illuminated open compartment (anxiogenic) for 30 s. The inter trial intervals were randomized from 2 to 5 minutes. Three high-resolution cameras operating at 30 Hz were used to record animal behavior. One camera was positioned above the FAST platform to capture movements from a top view. Two additional cameras were placed 15 cm away at the same height as the platform edges, oriented perpendicularly to each other to capture side views from orthogonal directions. This configuration ensured comprehensive coverage of animal movements across all axes. All cameras were synchronized using OBS Studio to ensure precise temporal alignment across recordings.

### Identification of ethologically defined behaviors

Animal behavior was recorded simultaneously from three synchronized cameras (top and two side views). For analysis, videos from all three perspectives were examined concurrently to ensure accurate identification of behavioral phenotypes. Behavioral scoring was performed manually by two independent experimenters who were blinded to the experimental conditions, and the results were compared to minimize observer bias. Six ethologically defined behaviors were identified and characterized: (1) grooming, repetitive cleaning or scratching movements of the face and body; (2) climbing, active rearing and grasping of the wall or edge; (3) walking, continuous locomotion across the platform; (4) protected stretch-attend posture (pSAP), body extension initiated from the enclosed or protected area toward the open area without full entry; (5) unprotected stretch-attend posture (uSAP), similar body extension occurring in open arms; and (6) head dips, forward lowering of the head over the edge of the open arm. The onset and offset of each behavioral event were manually annotated to generate time-stamped data for each behavioral phenotype. Each ethologically defined anxiety-related behavior was annotated separately in different video viewings, to minimize switching costs of manual scoring (*53, 54*). All analyses were conducted with experimenters blinded to group allocation.

### Spatial segregation of ethologically defined behaviors

To analyze the spatial distribution of each behavioral phenotype in the EPM, the open and closed arms were each divided into four equal segments, designated as C1, C2, C3, C4 (closed arms) and O1, O2, O3, O4 (open arms). This segmentation allowed for higher spatial resolution in determining the relationship between animal position and behavioral expression. For each behavioral event, the corresponding spatial zone was identified based on the animal’s location at the time of occurrence. The frequency and proportion of each behavioral phenotype within each zone were then calculated. All data from individual animals were subsequently pooled to generate the overall spatial distribution pattern of each behavioral phenotype across the EPM, which was further used for downstream analyses of spatial preference and correlations with neuronal activity.

### Ethologically defined behaviors transition analysis

To characterize the dynamic transitions among ethologically defined behaviors, we first extracted the start and stop time points for each behavioral event across the behavioral tests for all animals. Each behavior type (e.g., grooming, climbing, walking, pSAP, uSAP, and head dipping) was defined based on manually annotated video data. Using the timestamps of each behavioral phenotype, all events were sorted chronologically according to their start and stop times. Sequential or temporally overlapping events were identified as transitions from one phenotype to another.

For each animal, the number of transitions from one phenotype to another was counted, and transition probabilities were calculated by dividing transition count by the total number of transitions to all other behavioral states. A Markov chain analysis was then applied to compute the transition probabilities and visualize the overall behavioral state dynamics. The custom code used for this analysis is available on Figshare https://doi.org/10.6084/m9.figshare.30995806.

### Surgery

Mice were anesthetized with isoflurane (induction 3%, maintenance 1.5%) in oxygen at a flow rate of 1 L/min throughout the procedure. Core body temperature was kept at 37 °C by a feed-back controlled heating pad (Harvard Apparatus, Germany). Ophthalmic cream was applied to avoid eye drying. Local analgesia was applied by injecting a mixture of 2% of lidocaine (Streuli Pharma, Switzerland) and 0.5% bupivacaine (Aspen Pharma, Switzerland) subcutaneously under the scalp. Anesthesia depth was confirmed by detecting deep breathing and lack of toe pinch reflex. After mice were mounted onto a stereotaxic frame (David Kopf Instruments, USA), the scalp was incised and craniotomies were made over target regions. The surgical opening on skull is about Ø 0.1 mm for rabies tracing experiments, Ø 0.3 mm for optical fiber implantation and Ø 0.7 mm for gradient-index (GRIN) micro lens implantation. Mice were given additional analgesia for 3 days after operation (carprofen, 5 mg/kg subcutaneously). Implants e.g. optical fibers and GRIN lenses were secured to the skull using light-cured dental adhesive (Kerr, OptiBond Universal) and dental cement (Ivoclar, Tetric EvoFlow).

### Virus injection and GRIN lens implantation for miniscope

Conditional GCaMP6f viruses were injected into the mouse brain with the following coordinates: AP -3.28 mm, ML +3.45 mm, DV -4.0 mm relative to bregma. 250 nL viral solution was delivered via a glass micropipette (about Ø 20 μm at tip) attached by a tubing to a Picospritzer III microinjection system (Parker Hannifin Corporation) at a speed of 20 nL per minute. For labelling of pyramidal neurons, AAV1.CaMKII.GCaMP6f virus (Addgene #100834, titer 2.3×10^13^ vg/mL, dilution 1:3 for injection) was used in C57BL/6J mice. For labeling of PV, Sst or VIP interneurons, AAV5.Syn.Flex.GCaMP6f (Addgene #100833, titer 7×10^12^ vg/mL) was used in combination with each GABAergic Cre mouse lines.

After infusion of viruses, the glass micropipette was kept in place for 40 - 60 minutes to prevent viral solution backflow. Then the glass pipette was slowly retracted. The needle endomicroscope GRIN lens (Ø 0.6 mm) was slowly lowered to DV -4.0 mm at a speed of 0.5 mm per minute by using a leading 21 gauge needle attached to a custom-made stereotaxic guide enabling precise placement of the lens. The lens was fixed to the skull surface with light-cured dental adhesive and dental cement. The surface of the skull was made coarse by scratching with blade or gentle drilling to increase the adhesion with dental cement. Three skull screws were inserted around the implantation site and cemented together with the GRIN lens to ensure the implant’s stability. Additional cement was placed around the lens to form a well-shape cave to protect lens edge from damage.

From the third week post virus injection, GCaMP6f expression was inspected several times over consecutive weeks. Mice with good virus expression were fixed in a stereotaxic frame under isoflurane anesthesia to attach an aluminum baseplate for miniscope (UCLA, V3) above the GRIN lens. After finding the best field of view, the baseplate was cemented onto the skull and a plastic cap was used to protect the GRIN lens from dust. Mice wore a dummy miniscope for 2 weeks to adapt to the additional weight on head before behavioral tests and Ca^2+^ signal imaging.

### Virus injection and fiber implantation for optogenetic manipulation experiments

For bilateral optogenetic manipulation experiments, viruses encoding the inhibitory opsin eNpHR3.0 or excitatory opsin ChrimsonR, or a control tdTomato were injected at the following coordinates: AP -3.15 mm, ML ±3.0 mm, DV -4.5 mm relative to bregma. These coordinates are slightly different from the miniscope setting because the small size of optical fiber (Ø 0.2 mm) allowed us to penetrate deeper into the vH without too much damage to hippocampal structures. For the inhibition of pyramidal neurons, 400 nL AAV5-CaMKIIα-eNpHR3.0-mCherry virus (UNC Vector Core, titer 5.8×10^12^ vg/mL) was used in C57BL/6J mice. 400 nL AAV2-CaMKIIα-mCherry virus (UNC Vector Core, titer 4.7×10^12^ vg/mL) was used as control. For the inhibition of Sst and PV interneurons, 500nL AAV1-hEF1α-dlox-eNpHR3.0-iRFP-dlox virus (ETH Zurich Viral Vector Facility, v203, titer 3.8×10^12^ vg/mL) was used in either Sst.IRES.Cre or PV.IRES.Cre mouse line. 400 nL AAV8-EF1α1.1-Flex-tdTomato virus (UNC Vector Core, 4.5×10^12^ vg/mL) was used as control. After injection, the glass micropipette was left in place for another 20 minutes and then withdrawn slowly.

Optical fibers (200 μm core, 0.37 NA; Thorlabs) were cleaved to the appropriate length and secured to ceramic ferrules (Ø 230 μm bore, Senko) with tiny epoxy glue. After retracting glass micropipette, the optical fibers were attached into a stereotaxic cannula holder (Doric Lenses, Canada) and slowly inserted into the brain tissue at the same coordinate of virus injection.

### Virus injection and fiber and GRIN lens implantation for dual-color miniscope

For dual-color miniscope imaging experiments, the GRIN lens was implanted into the right hemisphere of the vH and the optical fiber on the left. The virus injection coordinates, fiber or GRIN lens implantation were the same as described in the above sections. For dual-color miniscope imaging of pyramidal neurons while inhibiting Sst or PV interneurons, a mixture of 100 nL AAV1.CaMKII.GCaMP6f and 400 nL AAV1-hEF1α-dlox-eNpHR3.0-iRFP-dlox was injected into the right hemisphere of the vH (AP -3.28 mm, ML +3.45 mm, DV -4.0 mm) and 500 nL AAV1-hEF1α-dlox-eNpHR3.0-iRFP-dlox into the left hemisphere of the vH (AP -3.15 mm, ML ± 3.0 mm, DV -4.5 mm). For controls, 400 nL AAV8-EF1α1.1-Flex-tdTomato was injected together with AAV1.CaMKII.GCaMP6f.

After virus injection, 2 skull screws were fixed at a position anterior to the bregma. An optical fiber was first implanted into the left hemisphere and secured with dental adhesive, and then the GRIN lens into the right hemisphere according to the procedure described in the above section. Dental cement was applied to secure screws, fiber and GRIN lens onto the skull. Mice were then returned to the animal facility and housed individually. 10 days of postoperative care and 3 days of analgesia were provided to these mice.

### Ca^2+^ imaging in freely behaving mice

Imaging sessions in freely-moving mice began 1 - 2 weeks after baseplating. Mice were briefly anesthetized (< 2 min) to attach the miniscope to the baseplate for each imaging day. Mice were allowed to recover from the brief anesthesia and habituate to the miniscope and behavior room for 60 minutes before the behavioral protocol started. Ca^2+^ imaging was performed using a miniscope (UCLA V3) or a custom-made dual-color miniscope for simultaneous Ca^2+^ imaging and optogenetic manipulation. The power of the blue laser used for GCaMP6f excitation (488 nm, Cobolt) was set to 1 mW at the tip of the miniscope objective. The power of the amber laser used for eNpHR3.0 and ChrimsonR excitation (594 nm, Cobolt) was set to 8-10 mW at the tip of the miniscope objective. The 488 nm laser was triggered by a Transistor-Transistor-Logic (TTL) signal from ANY-maze at the beginning of each recording session. The 594 nm laser was switched on according to optogenetic manipulation protocols. Ca^2+^ imaging videos were recorded at 20 Hz in uncompressed .avi format by using a data acquisition box which is triggered by an external TTL pulse from ANY-maze to synchronize Ca^2+^ imaging and behavioral tracking. The excitation power for GCaMP6f in pyramidal neurons was determined in prior tests based on the most optimal signal to noise ratio and was maintained throughout all the imaging sessions. Excitation power was sometimes slightly increased for PV interneurons imaging due to bleaching.

### Optogenetic manipulation

To optogenetically manipulate ventral hippocampal neuronal activity, a laser (Cobolt) generating 594 nm amber light was attached to an optical rotary joint (Doric Lenses) to support the unrestricted movement of mice during the behavioral tests. The optical rotary joint was connected to a light splitter (Doric Lenses) to allow bilateral light delivery to two patch cables (Doric Lenses) which were in turn connected to the implanted optical fibers through a ferrule-sleeve system (Senko, USA). Light illumination was automatically controlled by ANY-maze based on the animal’s body center position in the behavioral apparatus. Laser light was applied continuously at a power intensity of about 9-12 mW measured from the optical fiber tip. Before the beginning of the behavioral paradigm, mice were first connected to the patch cables for 30 minutes for habituation.

For the dual-color miniscope imaging, the laser source was divided into two patch cords by a light splitter, one for the optical fiber implanted into the left hemisphere, another one for dual-color miniscope into the right hemisphere. Because miniscope lost about 75% of light power through its optical pathways, the laser power for optical fiber was adjusted to similar level as at the tip of optical fiber by using an attenuating patch cord (Doric lenses). Ultimately, both brain hemispheres received approximately 9-12 mW laser light power.

During EPM tests, the optogenetic laser light was switched on whenever the mice body center was in the center zone or open arm during 121 s - 240 s, 361 s - 480 s, 601 s - 720 s in a 14-minutes long protocol.

### Immunohistochemistry

Mice were overdosed with a ketamine/xylazine cocktail and transcardially perfused with phosphate-buffered saline (PBS) followed by 4% formaldehyde. The brains were removed and fixed in 4% PFA for 24 hours at 4°C. Brains were sectioned coronally at a thickness of 50 μm using a vibratome (Leica Microsystems, VT1000S). The immunostaining was performed on free-floating brain slices. The sections were incubated in a blocking buffer containing 5% normal donkey serum (Jackson Immuno Research, Dianova) and 0.1% Triton-X 100 (Sigma, USA) in PBS for 1 hour to prevent nonspecific background staining. After blocking, the sections were incubated with the primary antibodies diluted in the blocking buffer. The following primary antibodies were used: anti-GFP polyclonal antibodies (Rockland 600-101-215, 1:2000; Invitrogen A11122, 1:2000), anti-Parvalbumin monoclonal antibody (Synaptic Systems, 195308, 1:1000), anti-Somatostatin monoclonal antibody (Invitrogen MA516987, 1:500; Millipore MAB354, 1:500), anti-RFP monoclonal antibody (Invitrogen MA515257, 1:1000). After overnight incubation with primary antibodies at 4°C, the sections were rinsed in PBS three times for 10 minutes each and incubated with secondary antibodies (2% BSA and 1% Triton-X 100 in PBS) for 2 hours in the dark at room temperature. The secondary antibodies were as follows: Goat anti-Rabbit IgG-conjugated Alexa Fluor 633 (A21070), Goat anti-Guinea Pig IgG-conjugated Alexa Fluor 633 (A21105), Donkey anti-Rabbit IgG-conjugated Alexa Fluor 405 (A48258), Donkey anti-Rat IgG-conjugated Alexa Fluor 405 (A48268), Donkey anti-Rat IgG-conjugated Alexa Fluor 594 (A21209), Donkey anti-Rabbit IgG-conjugated Alexa Fluor 594 (A21207), Donkey anti-Rabbit IgG-conjugated Alexa Fluor 488 (A21206), Goat anti-Guinea Pig IgG-conjugated Alexa Fluor 594 (A11076), all purchased from Invitrogen used in dilution of 1:1000. Afterwards, the sections were washed three times in PBS for 10 minutes, transferred to Superfrost Plus charged glass slides, and mounted in Vectashield mounting medium (Vector Laboratories, USA).

### Ca^2+^ imaging data extraction

Only mice with verified viral expression and GRIN lens placements in the target sites were included in the analyses. Ca^2+^ imaging videos were analyzed using a custom-made Matlab code(*55*). Videos from multiple sessions were concatenated and downsampled by a binning factor of 4 resulting in a frame rate of 5 Hz, and lateral brain movements were motion-corrected using the Turboreg algorithm. Fluorescent traces were extracted by applying automatically detected individual cell filters based on combined principal and individual component analysis (PCA/ICA). This approach combines spatial and temporal statistics and precedes ICA and PCA, to reduce data dimensionality and to support ICA in finding global optima. The overall procedure is proved to be effective from grounded hypotheses: cellular signals are mathematically separable into products of paired spatial and temporal components; signals from different cells are statistically independent; and cells’ spatial filters and temporal signals have skewed distributions. In brief, this automated sorting procedure combines ICA and image segmentation for extracting cells’ locations and their dynamics with minimal human supervision. To control for non-inclusion of split neurons in our analyses, we identified pairs of neurons with highly correlated activity (Pearson correlation > 0.7) that were spatially close (centroid distance < 20 pixels) and excluded one of the neurons for each pair. Identified putative neurons were then sorted via visual inspection to select neurons with appropriate somatic morphology and Ca^2+^ dynamics.

### Neuronal activity analysis

Relative changes in Ca^2+^ fluorescence (F) were calculated using the formula: ΔF/F_0_ = (F – F_0_) / F_0_ (F_0_ = the fluorescence intensity over the entire trace) and used for all the analyses of Ca^2+^ activity. Ca^2+^ transient level was used as a proxy of neuronal activity. The neuronal activity level was presented as integral of area under the curve of Ca^2+^ transient normalized by the time of the corresponding period (AUC/s). Ca^2+^ signal that was 2 times higher than the standard deviation (SD) of the entire trace was considered as a relevant neuronal activity. All analyses were done on z-scored traces.

### Correlation of neuronal activity with initiation of behavioral phenotypes

To analyze the neuronal activity patterns associated with the initiation and termination of behavioral phenotypes, calcium signals were aligned to the start or stop time points of each behavioral event. For each ethologically defined begavior, calcium traces were extracted and averaged within a 1.5-second window before and after the onset of the behavior. This alignment enabled the visualization of neuronal activity dynamics surrounding behavioral transitions. The custom MATLAB code used for this analysis and plotting is available on Figshare (https://doi.org/10.6084/m9.figshare.30995806).

## Statistics and Reproducibility

Analyses were performed using custom scripts written in MATLAB (MathWorks). Statistics were performed using GraphPad Prism 9. All datasets were tested for normality using the Kolmogorov-Smirnov test or Shapiro-Wilk test based on sample size. Normally distributed data undergo parametric tests, otherwise nonparametric tests were applied. All null hypothesis tests were two-sided. Analyses of variance (ANOVAs) were followed by post hoc tests if a main effect or interaction was observed. Box and whisker plots show median and interquartile range (minima, 25^th^ and 75^th^ percentile, and maxima). Bar graphs show data as means ± SEM. Asterisks in the figures represent *P* values corresponding to the following thresholds: \**P* < 0.05; \*\**P* < 0.01; \*\*\**P* < 0.001.

## Data Availability

Source data are provided with this paper. The original data used in this study are available in Figshare repository under open access licence CC BY 4.0. https://doi.org/10.6084/m9.figshare.30995806. Additional data relating to this paper are available upon request, because of the size of the calcium imaging and animal behavioral tracking data is too large to be deposited online.

## Code Availability Statement

The Matlab code for AUC analysis, behavior phenotype transition analysis, initiation and termination of behavior phenotypes used in this study is deposited in Figshare, doi: https://doi.org/10.6084/m9.figshare.30995806.

## Acknowledgements

This study was supported by a European Research Council starting grant 716761 (SC), Swiss National Science Foundation professorship grant 206129 (SC), Novartis Foundation for medical-biological Research grant 24C258 (SC), Swiss National Science Foundation Flexibility grant (KL), Novartis Foundation for medical-biological Research grant 24A021 (KL).

## Author contributions

Conceptualization: KL, SC; Methodology: KL; Investigation: KL; Visualization: KL; Funding acquisition: KL, SC; Project administration: KL; Data acquisition: KL; Data analysis: KL, MK; Supervision: SC; Writing: KL, SC, MK.

## Supplementary figures

**S1.**
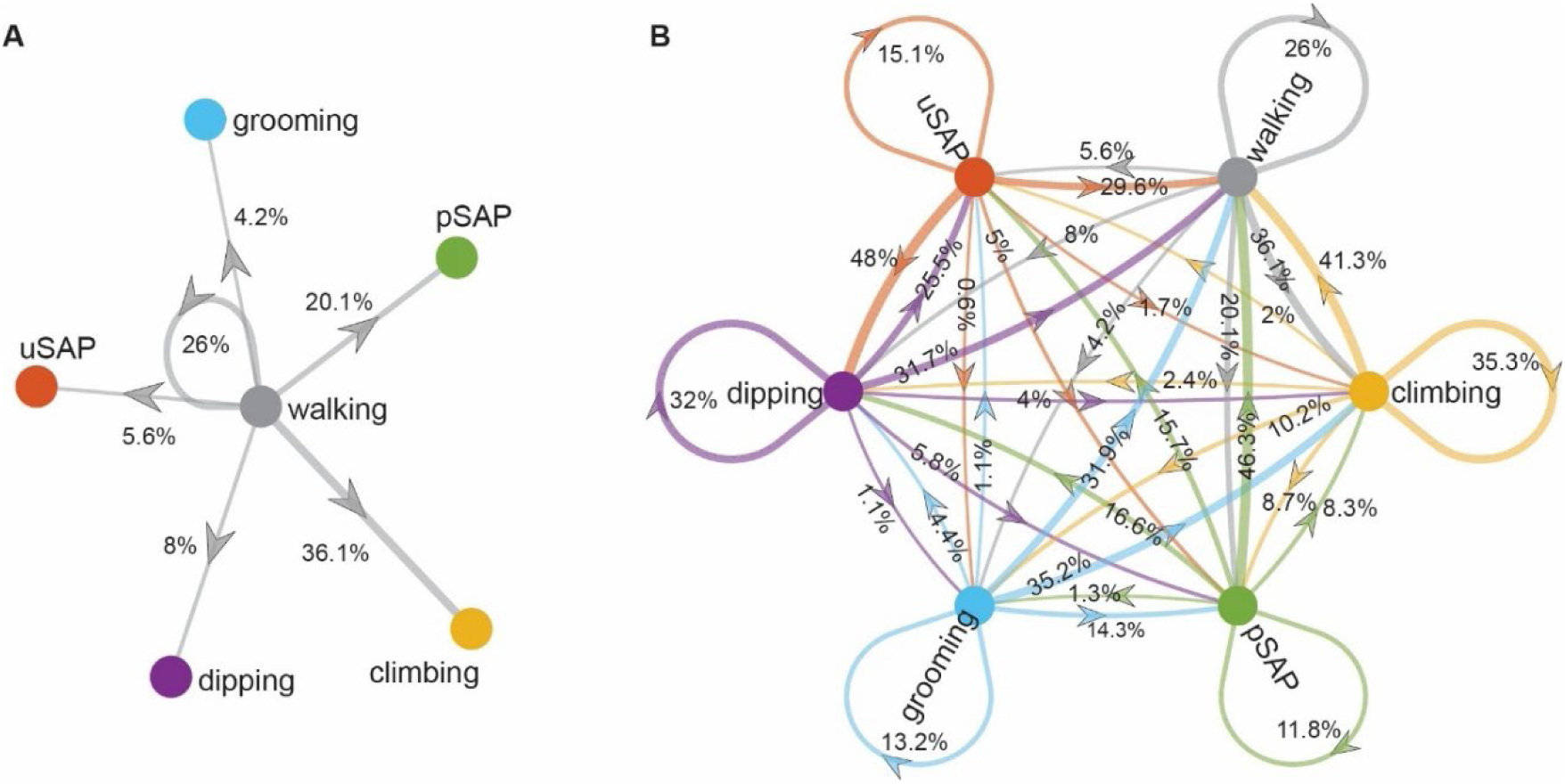
Markov chain analysis of transition probability among ethologically defined behaviors. (**A**) Transition probabilities from walking to other ethologically defined behaviors. (**B**) Transition probabilities among all ethologically defined behaviors.

**S2.**
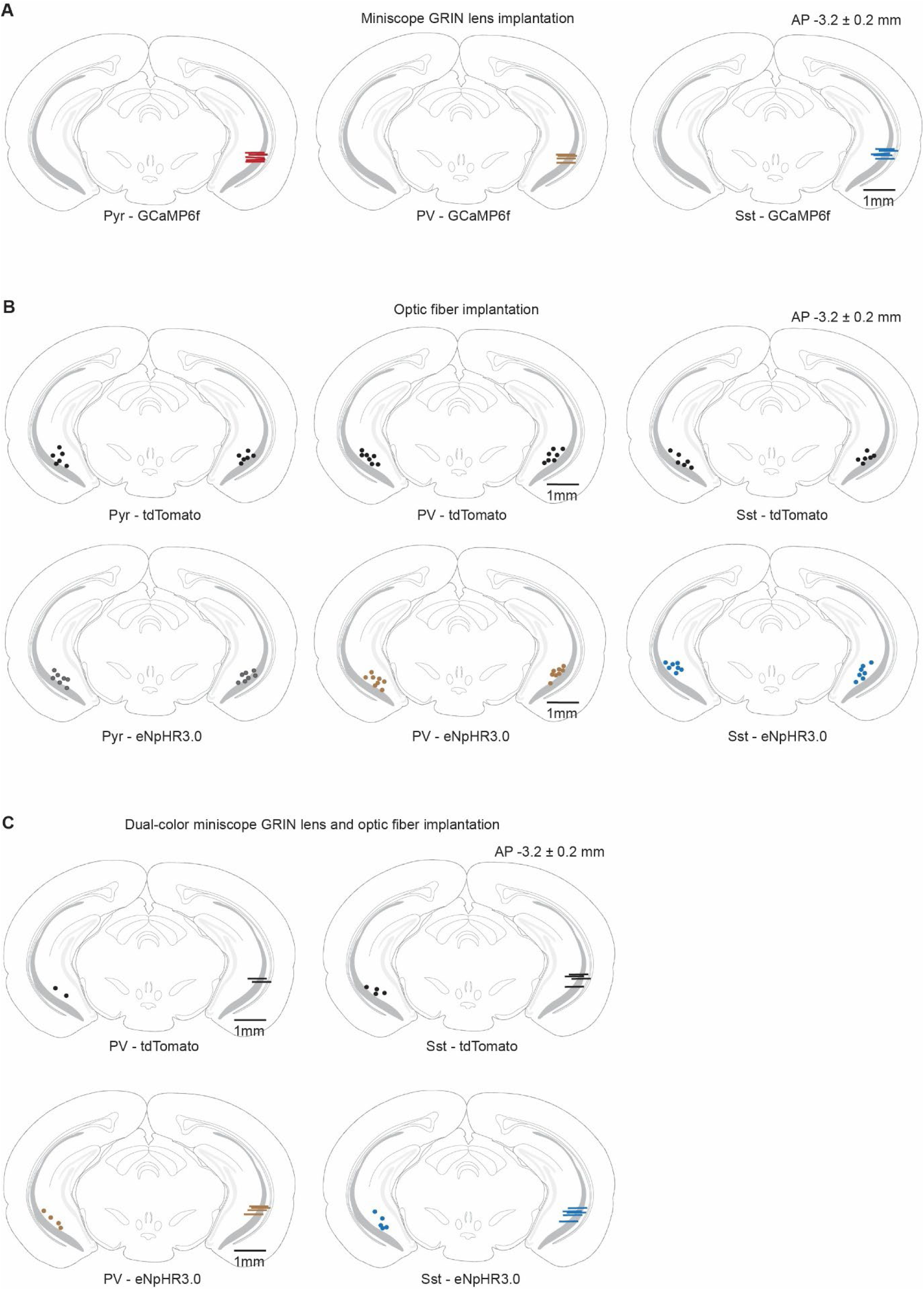
Optical fiber and GRIN lens implantation. Schematic illustrating reconstructed implantation sites of optical fiber (**A**) for optogenetic manipulation, GRIN lenses (**B**) for miniscope imaging, and optical fiber and GRIN lens (**C**) for dual-color miniscope. Each dot or line represents one mouse.

**S3.**
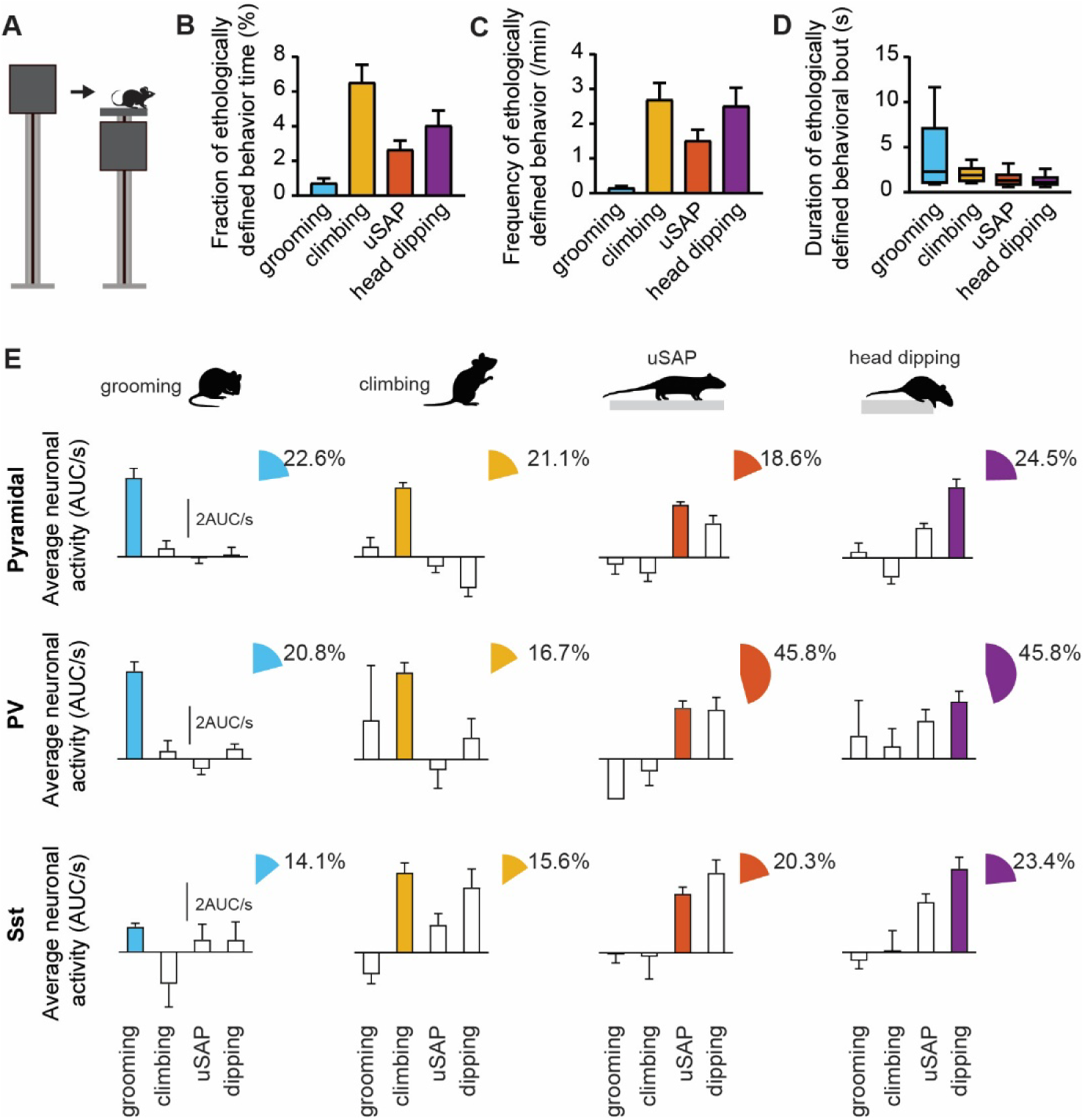
Ethologically defined behaviors selectively recruit subclasses of vH neurons on FAST. **(A)** Schematic of forced anxiety shifting task (FAST) protocol. **(B)** Fraction of time of each ethologically defined behaviors during test. (**C**) Frequency of each ethologically defined behavior per mouse. (**D**) Duration of ethologically defined behavioral bouts. n = 8 mice. **(E)** Averaged neuronal activity and fraction of ethologically defined behaviors with selectively recruited neurons on FAST. n = 204 neurons from 5 C57BL/6 mice; n = 25 neurons from 2 PV.Cre mice; n = 64 neurons from 5 Sst.Cre mice.

